# Disrupted Development of the mPFC-Thalamic Circuit in the Mouse Shank3^−/−^ model of Autism

**DOI:** 10.1101/2024.09.09.612160

**Authors:** Gabrielle Devienne, Gil Vantomme, John R Huguenard

## Abstract

Autism Spectrum Disorders (ASDs) are a group of neurodevelopmental disorders with heterogeneous causes, and are characterized by communication deficits, impaired social interactions, and repetitive behaviors. Despite numerous studies in mouse models focused on pathophysiological circuit mechanisms of ASD in mature animals, little is known regarding ASD onset and its evolution through development. The medial prefrontal cortex (mPFC) is crucial for higher-order cognitive functions and social behavior, thus key to understanding ASD pathology. To explore early developmental disruptions in the mPFC, we used the *Shank3* knockout (Shank3^−/−^) mouse model. SHANK3 is crucial for glutamatergic synapse maturation, and the Shank3^−/−^ mouse has been well-characterized for displaying ASD-related behavioral phenotypes. We investigated network, cellular, and synaptic changes within the mPFC and in its projections to the mediodorsal thalamus (MD) at two developmental stages, preweaning (P14) and adulthood (>P55). Our findings reveal early synaptic deficits at P14 both within the mPFC and in its projections to the MD accompanied by alterations in mPFC network activity and reduced excitability of excitatory neurons, overall suggesting hypofunction. Interestingly, behavioral deficits were already detectable by P11, preceding the observation of synaptic changes at P14. By adulthood, these early synaptic and cellular alterations progressed to global dysfunction, characterized by mPFC network hyperfunction and layer 5 pyramidal cell hyperexcitability, accompanied by augmented glutamatergic signaling to MD with enhanced action potential production. These results suggest that early synaptic changes may precede and interact with behavioral deficits, potentially leading to compensatory mechanisms that contribute to more pronounced mPFC dysfunction later in development. This study highlights the complex dynamic progression of mPFC deficits in ASD and emphasizes the potential impact of targeting early synaptic alterations to mitigate later behavioral and cognitive deficits.

## Introduction

Autism Spectrum Disorders (ASDs) refer to a group of neurodevelopmental disorders characterized by communication deficits, impaired social interaction, and repetitive behaviors. ASD arises from complex etiologies including genetic causes and prenatal insults, making it difficult to precisely determine its onset across individuals. In humans, the behavioral deficits related to ASD are generally first observed around the second year of life(1,2), a period associated with rapid maturation of the medial prefrontal cortex (mPFC)(3). Abnormal mPFC development is hypothesized to contribute to ASD symptoms(4–6), given its critical role in regulating social behavior(5,7–9). In mouse, communication deficits can be perceived in the early postnatal periods (pre-weaning; <P21)(10,11) a critical window for morphological and functional mPFC development (12). It is around the same period that mPFC develops long range connections with subcortical areas(12,13) including reciprocal connections with the mediodorsal thalamus (MD)(14).

SHANK3, a scaffolding protein critical for glutamatergic synapse maturation(15), is a leading genetic cause of ASD(16,17). The Shank3^−/−^ mouse model(11) recapitulates many ASD-related phenotypes, including social deficits and communication impairments. Interestingly, SHANK3 protein levels naturally decline around P14 during normal development(18), coinciding with a reduction in connectivity between the mPFC and the MD(19). This developmental window offers a unique opportunity to investigate how early synaptic changes might predispose the mPFC-MD circuit to later dysfunction. By studying the Shank3^−/−^ model at both P14 and adulthood (>P55), we aim to identify whether early synaptic alterations set the stage for later deficits in network and behavioral function.

To investigate mPFC disruptions in Shank3^−/−^ mice, we compared network, cellular, and synaptic properties at P14 and adulthood (>P55). Behavioral assessments confirmed communication deficits as early as P11(11) as well as social deficits in adulthood(11,20), emphasizing the early onset and persistence of ASD-related symptoms. Electrophysiological methods, including local field potential (LFP) and whole-cell patch-clamp recordings, were used to examine mPFC activity. We also tested mPFC-MD circuit function using optogenetic stimulation of mPFC axons and intracellular recordings in MD cells.

Our findings reveal a dynamic progression of mPFC deficits in Shank3^−/−^ mice. At P14, mPFC hypoexcitability was observed in upper layer pyramidal cells, with synaptic excitatory-inhibitory imbalance in layer 5. Despite an overall mPFC network hypoexcitability, optogenetic stimulation of mPFC-MD projections revealed exuberant and long-lasting synaptic excitation in MD cells, an early sign of circuit hyperexcitability. In adulthood, pronounced mPFC network hyperactivity emerged, particularly in layer 5 pyramidal cells that project to the MD. Long lasting excitatory synaptic responses in MD cells were persistently augmented in Shank3^−/−^ mice, indicating long-term disruptions in mPFC-MD connectivity.

These results show that early synaptic changes in Shank3^−/−^ mice precede pronounced network dysfunction in adulthood. Mixed findings at P14 suggest compensatory mechanisms may temporarily mask mPFC dysfunction, highlighting critical periods of vulnerability in ASD and the potential for early interventions to mitigate long-term deficits.

## Results

### Early emergence of social behavioral deficits in Shank3^−/−^ mice

To assess the development of potential SHANK3-related behavioral deficits, we tested WT and Shank3^−/−^ mice at two distinct developmental stages: pups (P11) and adults (>P55). Given the limited mobility of pre-weaning pups, ultrasonic vocalizations (USV) have be used as a proxy of social behavior(21). Upon social isolation Shank3^−/−^ P11 pups produced many fewer calls compared to their wild type (WT) littermates (Figure 1 A-D; mean ± sem; WT: 253 ± 58 calls; Shank3^−/−^: 61 ± 30 calls; testing period 8 minutes; Mann-Whitney, p=0.01). Consistent with social deficits persisting into adulthood, mature Shank3^−/−^ mice failed to show a social preference in a social interaction test, showing similar times with either a novel inanimate object or a novel conspecific mouse. In contrast, adult WT mice showed a strong social preference (Figure 1 E-G; interaction time, mean ± sem; WT: mouse, 33.1 ± 4.95, object, 15.8 ± 1.95 %, paired t-test, p= 0.02; Shank3^−/−^: mouse, 14.6 ± 5.62, object: 15.8 ± 5.91%, paired t-test, p= 0.89). These results align with previous literature on ASD mouse models(22–24), including the complete Shank3 knockout used here(11). We therefore confirm that behavioral deficits in this model begin as early as P11, which correlates with a period of critical development of the corticothalamic network connecting the medial prefrontal cortex (mPFC) to the mediodorsal thalamus (MD) (12,14), an important circuit related to social behavior(25).

**Figure 1.**
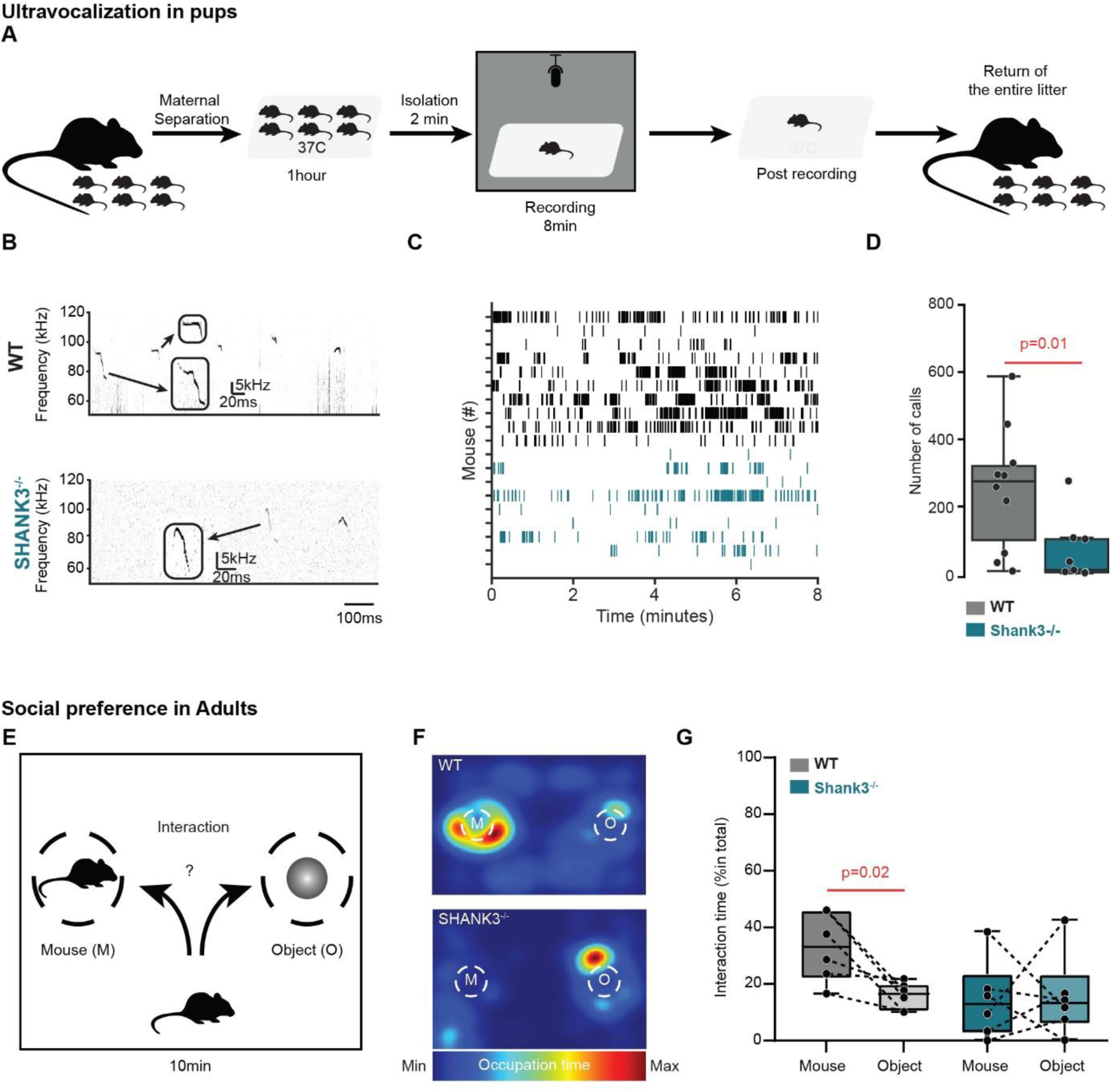
Shank3^−/−^ mice display deficits in communication in early life and in social behavior in adulthood. (A) Schematic diagram of isolation and ultrasonic vocalization protocol performed in Shank3 WT and Shank3^−/−^ P11 littermates. (B) Examples of 1 second long spectra (bottom images, in kHz) from USV recordings in WT and Shank3^−/−^ mice. Insets show examples of individual syllables expanded as indicated by arrows in the original recordings. Left: WT (black); Right: Shank3 (blue). (C) Raster plot showing individual mouse ultrasonic vocalization (USV) calls during 8-minute recordings, with WT mice represented in black and Shank3^−/−^ in blue (D) Number of calls for WT (grey) and Shank3^−/−^ (blue). (E) Schematic diagram of social preference test performed in WT and Shank3^−/−^ adults. (F) Example of occupation time heat maps for WT (top) and Shank3^−/−^ (bottom) adults. (G) Percentage of the interaction time with a never-met conspecific (darker colors) or a novel inanimate object (lighter colors) for WT (grey) and Shank3^−/−^ (blue).

### Developmental progression of hyperexcitability in the mPFC network

To investigate potential SHANK3-related disruptions in mPFC development, we first employed an *in vitro* extracellular recording approach to evaluate network excitability. We obtained coronal brain slices containing the mPFC from Shank3^−/−^ mice and WT littermates at different developmental stages, including P11, P14, and adulthood (Figure 2 and Figure S1). Using a 16-shank linear silicon probe array positioned normal to the pial surface of mPFC, we recorded network activity in the form of local-field potentials (LFPs). Local electrical stimulation of deep layers evoked multicomponent LFPs that could be observed across all cortical layers, demonstrating engagement of intra-cortical circuits (Figure 2A). To localize the LFP signals to different cortical layers, we calculated the current source density(26) (CSD) as the second spatial derivative of the LFP (Figure 2B and C). This allowed us to compare mPFC network differences between Shank3^−/−^ mice and WT littermates. We observed matched pairs of sinks and sources in the CSD profiles following local electrical stimulation in the deep layers. Sinks are negative CSD signals generally attributed to the movement of cations from the extracellular space into the cells during synaptic excitation. Given conservation of charge, such ionic current flow must be matched with a paired outward flow of cations (sources) from these cells at a distance from the original sinks. Synaptic inhibition has the opposite sinks/source polarity as it is mainly associated with anionic, rather than cationic, current flow. Accordingly, sources/sinks cannot unambiguously be associated with either excitation or inhibition. However given that the majority of synapses in neocortical circuits are excitatory(27–29), sinks will generally be dominated by excitatory synaptic signals, and sources with their return path. We applied a subtractive pharmacological approach to isolate components of the CSD signal, to reveal distinct pre- and post-synaptic components, with the post-synaptic components essentially blocked by DNQX and CPPene, AMPA and NMDA receptor antagonists (Figure S1A2, B), and presynaptic components blocked by TTX.

**Figure 2.**
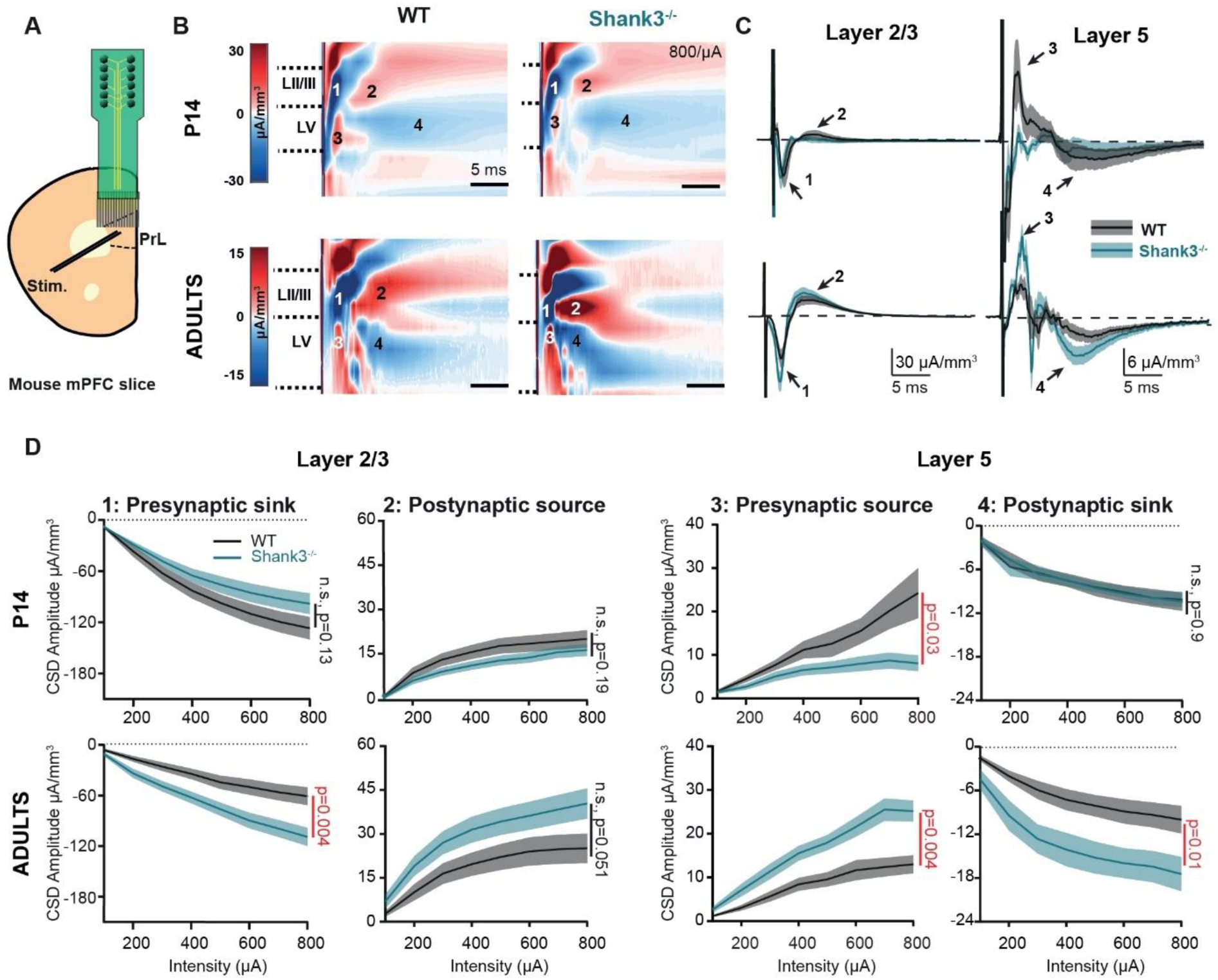
Shank3^−/−^ mice display mPFC hypofunction in early life and hyperfunction in adulthood. (A) Schematic of LFP recording across all mPFC layers in response to electrical stimulation delivered to deep layers. (B) Average current source density (CSD) response displayed as a heat map of mPFC network activity in response to a near maximal (800µA) stimulation in WT (left) and Shank3^−/−^ (right) at P14 (top row; WT= 3 mice, 10 slices; Shank3^−/−^ = 3 mice, 11 slices) and adults ( >P55, bottom row; WT= 5 mice, 12 slices; Shank3^−/−^= 5 mice, 13 slices). (C) Average CSD response of the L2/3 presynaptic sink (arrow 1, numbers correspond to those in B), L2/3 postsynaptic source (arrow 2), L5 presynaptic source (arrow 3) and L5 postsynaptic sink (arrow 4) in P14 (upper) and adult (lower) slices. Traces represent mean (solid line) ± sem (shaded line). (D) Input/ Output curve of CSD peak amplitude as a function stimuli intensity for P14 (top row) and adults (bottom row); WT (grey) and Shank3^−/−^ (blue).

We initially assessed mPFC excitability at the two stages that we had previously characterized behaviorally, P11 and adult. At P11 we observed LFP and CSD components generally comparable to what was observed in adult mice, although with somewhat simpler features especially noted as more monophasic responses, consistent with the immature state of the network at P11 (Figure 2A, vs. Figure S1C). We did not detect any differences between Shank3^−/−^ mice and their WT littermates at this stage. We then selected P14 as the next developmental time point. This is a developmental stage marked by significant changes in SHANK3 protein levels(18). The CSD signal at P14 showed sinks and sources across layers that were more complex (multiphasic) than at P11, and more similar to the CSD responses in adults. At P14, both the previously identified postsynaptic sinks and sources (Figure S1B) were similar between Shank3^−/−^ mice and WT littermates. However, we observed a significant reduction of the presynaptic source located in layer 5 (Figure 2C, component 3),with little to no reduction in presynaptic sink in layer 2/3 (component 1) in Shank3^−/−^ mice compared to WT littermates (Figure 2CD P14; two-way ANOVA with repeated measures: genotype p=0.03, stimulation p<0.0001, interaction p<0.0001 for the layer 5 presynaptic source). This reduction in CSD signal may reflect a hypofunction of mPFC at this stage of development, in particular of the presynaptic recruitment of the cortical network. By contrast, at adulthood, mPFC slices from Shank3^−/−^ mice showed an overall *increase* in sinks and sources across layers (components 1-4, Figure 2B-D). Closer inspection revealed that this overall increase in CSD signal in adult is mediated by an increase in the peak amplitude of the layer 2/3 presynaptic sink and the layer 5 presynaptic source and postsynaptic sink (Figure 2CD Adults; 2-way ANOVA with repeated measure, layer 2/3 presynaptic sink: genotype p=0.004, stimulation p<0.0001, interaction p<0.0001; layer 5 presynaptic source: genotype p=0.001, stimulation p<0.0001, interaction p<0.0001, layer 5 postsynaptic sink: genotype p=0.01, stimulation p<0.0001, interaction p=0.04). There is also a trend for an increase in the peak amplitude of layer 2/3 postsynaptic source in adult Shank3^−/−^ mice (Figure 2D Adults; 2-way ANOVA with repeated measure, layer 2/3 postsynaptic source: genotype p=0.051, stimulation p<0.0001, interaction p=0.08). Altogether, these data demonstrate the emergence of a hyperfunctional mPFC network between P14 and adulthood (>P55) in the Shank3^−/−^ mouse model, characterized by early-stage hypoexcitability at P14 followed by hyperexcitability in adulthood, reflecting a developmental shift in network dynamics.

### Developmental emergence of hyperexcitability of layer 5 mPFC pyramidal cells

Our *in vitro* LFP experiments revealed increased mPFC network activity in Shank3^−/−^ adult mice, evidenced in both pre- and postsynaptic signals. These results suggest potential alterations in the properties of local mPFC cells and/or in the synaptic connections within the mPFC. Excitatory pyramidal neurons in the mPFC are crucial for regulating cognitive and social behavior(30). Layer 2/3 of the mPFC receives diverse long-range inputs, including notable projections from the MD thalamus(31). Conversely, layer 5 of the mPFC primarily projects to the MD thalamus, although not exclusively(12,32,33) as it can target other subcortical areas along with the contralateral mPFC.

To investigate changes in cellular excitability, we recorded mPFC layer 2/3 and layer 5 pyramidal cells at P14 and in adulthood (>P55). Consistent with decreased excitability observed in the LFP (Fig 2D), early in development L2/3 pyramidal cells revealed hypoexcitability, in particular with a hyperpolarized resting membrane potential (Figure 3A; RMP, P14; mean ± sem; WT: −58.6 ± 1.3 vs Shank3^−/−^: −62.8 ± 1.3; p=0.03, Welsh t-test) which is not observed in adults. In addition, we found that these cells fired significantly fewer action potentials in response to depolarizing current steps (Figure 3B; 2-way ANOVA; genotype p=0.0002, current injection p<0.0001, interaction p<0.0001). In adulthood this layer 2/3 hypoexcitability was no longer evident (Figure 3C).

**Figure 3.**
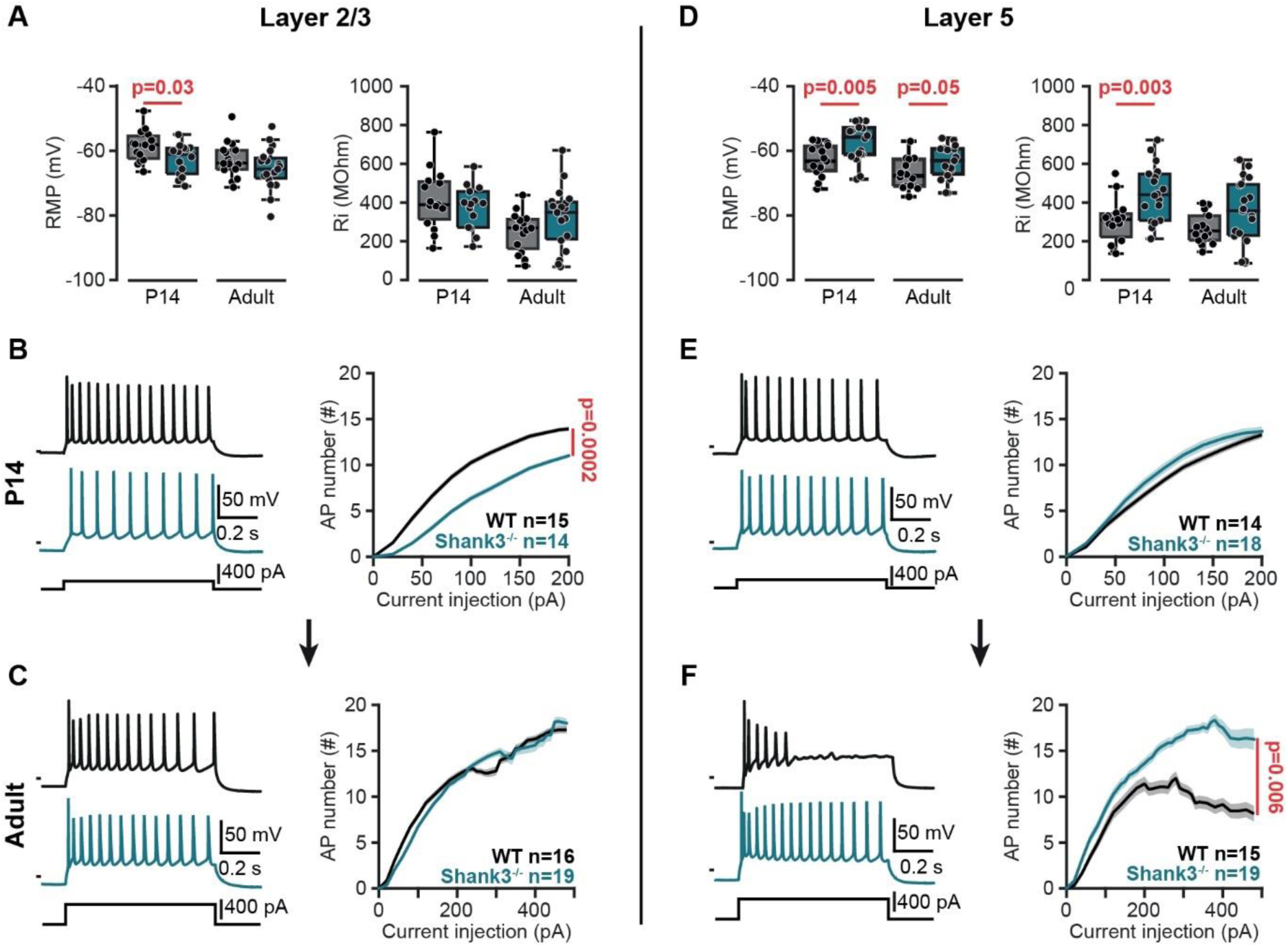
Shank3^−/−^ mice display early electrophysiological hypoexcitability in mPFC layer 2/3 that develops into hyperexcitability of layer 5 in adulthood. (A) Resting membrane potential (RMP, left) and input resistance (Rm, right) of layer 2/3 pyramidal neurons from P14 and adult WT (grey) and Shank3^−/−^ (blue) mice. (B) Left, representative action potential trains in layer 2/3 mPFC pyramidal cells in response to 100 pA, 1s current injection at P14. Right, number of APs in layer 2/3 pyramidal cells at P14. Traces represent mean ± sem. (C) Same as (B) in adult mice. (D) Resting membrane potential (RMP, left) and input resistance (Rm, right) of layer 5 pyramidal neurons from P14 and adult WT (grey) and Shank3^−/−^ (blue) mice. (E) Left, representative action potential responses of layer 5 mPFC pyramidal cells at P14 (400 pA stimulus). Right, number of APs in layer 5 pyramidal cells at P14. Traces represent mean ± SEM. (F) Same as (E) in adult mice.

Various ASD mouse models have been shown to display hyperexcitability of adult mPFC layer 5 pyramidal cells(5,34,35). Consistent with this, we find that in Shank3^−/−^ mice layer 5 pyramidal cells have a depolarized RMP (Figure 3D; RMP; Adults; mean ± sem; WT: −67 ± 1.3 vs Shank3^−/−^: −63.4 ± 1.2, p=0.048, Welsh t-test). Similar results were observed at P14 (Figure 3D; RMP; P14; mean ± sem; WT: −63 ± 1.3 vs Shank3^−/−^: −57.4 ± 1.4, p=0.005, Welsh t-test). This early sign of hyperexcitability is also reflected by a higher input resistance (Figure 3D; RMP; P14; mean ± sem; WT: 304 ± 29 vs Shank3^−/−^: 448 ± 33, p=0.003, Welsh t-test). At P14, layer 5 pyramidal cells displayed similar action potential numbers in response to a range of depolarizing current injections in Shank3^−/−^ and WT littermates (Figure 3E). Interestingly, in this assay Shank3^−/−^ adult layer 5 pyramidal cells display higher number of action potentials (Figure 3F; 2-way ANOVA; genotype p= 0.0058, current injection p<0.0001, interaction p<0.0001). In addition, these cells also display an increased hyperpolarization induced membrane potential sag compared to their WT littermates (Figure S2 B5, p=0.04, Mann Whitney test). We did not observe any alterations in individual action potential properties at any age or in any layer (Figure S2).

Biocytin was included in our intracellular recording solution, which allowed us to recover the cellular morphology of a subset of mPFC layer 5 pyramidal cells in both WT and Shank3^−/−^ at P14 and in adults. A Scholl analysis at both ages revealed an increase in dendritic complexity in Shank3^−/−^ cells (Figure S3B; 2-way ANOVA, P14, genotype p=<0.0001, distance p=<0.0001, interaction p=<0.0001; Adults, genotype p=<0.0001, distance p=<0.0001, interaction p=<0.0001). Shank3^−/−^ neurons had more arborizations near the soma and more branches overall compared to WT, while the average neurite length was similar (Figure S3C, D, E). It has been described that layer 5 pyramidal cells can be split in at least two different subtypes: subcortically projecting, thick-tufted neurons that display a marked H-current and related hyperpolarization induced membrane potential sag, and callosal projecting, thin-tufted neurons that display an absence in H-current(36,37). To exclude the possibility that these findings represent differences in the proportions of recorded cell subtypes in Shank3^−/−^ vs WT animals, we measured apical dendritic shaft width in WT and Shank3^−/−^, and these did not show any differences (Figure S3F) suggesting that our morphological analysis was performed in comparable mPFC neuron populations, such that the differences in cell morphology we describe are unlikely to be explained by differences in proportion of subtypes in WT vs. Shank3^−/−^ mice. Overall, WT/ Shank3^−/−^ mPFC layer 5 pyramidal neuron increases in excitability and dendritic complexity are already present at P14 and further develop into adulthood.

In summary, our whole-cell patch-clamp recordings revealed signs of hypoexcitability in mPFC layer 2/3 pyramidal neurons in juvenile Shank3^−/−^ mice. By contrast, at this stage, we already see early signs of hyperexcitability in layer 5 pyramidal cells which persist with greater functional consequences in adults.

### Decreased E-I ratio observed at P14 and in the adult Shank3^−/−^ mice

A major goal in ASD research has been to find convergent mechanisms in different genetic mouse models. One of the main hypotheses, as proposed by Rubenstein and Merzenich(38), is linking genetic models of autism through a common mechanism of increase in excitatory-inhibitory synaptic ratio. We used whole-cell voltage-clamp recordings to examine the synaptic responses in mPFC layer 2/3 and layer 5 pyramidal cells. Synaptic responses were evoked using an electrical bipolar stimulating electrode placed in upper layer 2/3. Excitatory post-synaptic currents (EPSCs) were recorded at V_m_ of −70mV and inhibitory postsynaptic currents (IPSCs) at V_m_ of +10mV (Figure 4A). The excitatory-inhibitory ratio in Layer 2/3 mPFC pyramidal cells were not different between WT and Shank3^−/−^ at either P14 or in adults (Figure S4B). By contrast, Shank3^−/−^ mPFC layer 5 pyramidal cells displayed a reduced E-I ratio already evident at P14 (Figure 4B-C; mean ± sem; P14 WT: 0.98 ±0.21, P14 Shank3^−/−^: 0.4 ± 0.1, Mann Whitney test, p=0.04) and reduction continued into adulthood (Figure 4B-C; mean ± sem; WT: 1.1 ± 0.3, Shank3^−/−^: 0.5 ± 0.1, Mann Whitney test, p=0.03). These data, in contrast to previous observations made in other ASD models, suggest that the imbalance of the E-I ratio in Shank3^−/−^ mice is an actual reduction in E-I ratio and is restricted to inputs to layer 5 pyramidal neurons of the mPFC. Interestingly, this imbalance is observed as early as P14, indicating that these alterations in synaptic function occur during early developmental stages, potentially contributing to the onset and progression of ASD in Shank3^−/−^ mice.

**Figure 4.**
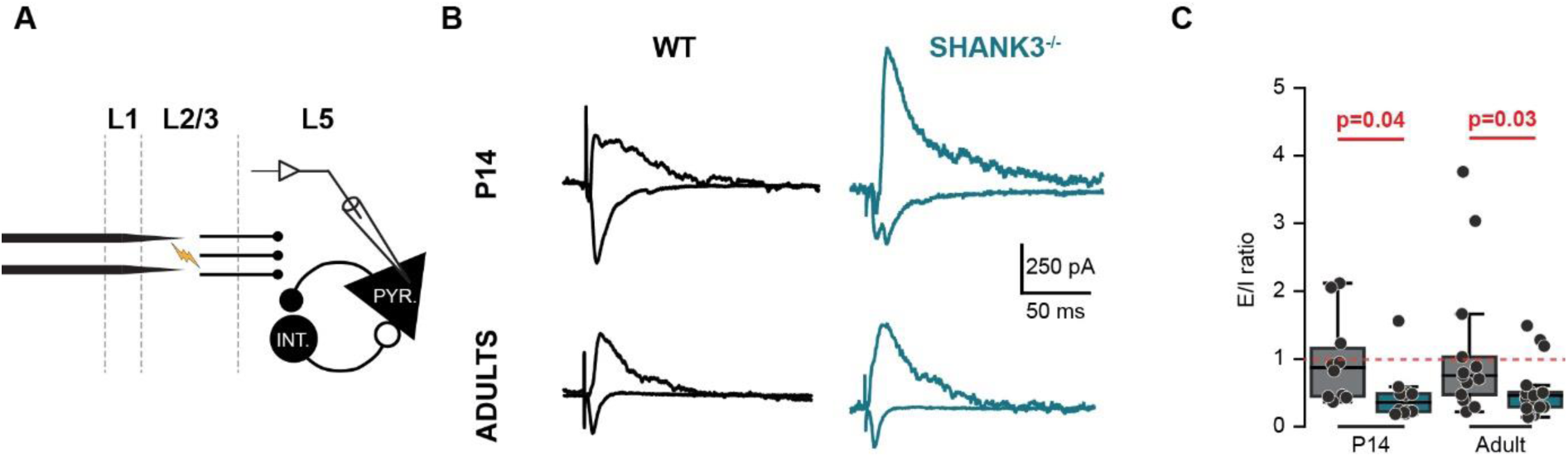
Excitatory/inhibitory synaptic current ratio is reduced in Shank3^−/−^ mPFC L5 pyramidal cells at both P14 and in adults. (A) Experimental design to measure EPSCs (at −70 mV) and IPSCs (at +10 mV) in mPFC slices, with electrical stimulation of layer 2/3. (B) Representative Excitatory and Inhibitory synaptic current responses elicited in mPFC layer 5 pyramidal cells by an electrical stimulation in layer 2/3 of P14 (top) and adult (bottom) WT (grey) and Shank3^−/−^ (blue) mice. (C) E/I ratio at P14 (left panel; WT (grey), 4 mice, 10 cells; Shank3^−/−^ (blue), 5 mice, 12 cells) and adulthood (right panel; WT (grey), 7 mice, 11 cells; Shank3^−/−^ (blue), 7 mice, 15 cells).

### Altered mPFC-MD thalamus synaptic function in Shank3^−/−^ mice

As outlined above, one of the major projections of mPFC layer 5 pyramidal cells is the MD thalamus. To evaluate whether the changes we observed in mPFC were associated with and potentially causative of effects on downstream targets, we injected mPFC with one of two Channelrhodopsin2 expressing viruses AVVDJ for P0 injections (Figure S6) and AAV1 CaMKIIa-ChR2-eYFP for adults, followed by in vivo incubation for several weeks. We then recorded synaptic responses in MD cells (Figure 5A Figure S6). Shank3^−/−^ MD neurons showed passive properties that were indistinguishable from controls (Figure 5A1-8), suggesting that the changes in intrinsic excitability we had observed were restricted to cortical locations. Using 1Hz blue light stimulation, we reliably evoked optogenetic monosynaptic excitatory synaptic currents (oEPSCs) as observed with a short and fixed latency (in ms; mean ± sem; P14: WT=4.3 ± 1.1; Shank3^−/−^ =4.9 ± 1; Welch’s t-test, p=0.1; adult: WT=2.6 ± 0.1; Shank3^−/−^ = 2.9 ± 0.1; Welch’s t-test, p=0.1). oEPSCs showed similar rise time between WT and Shank3^−/−^ (in ms; mean ± sem; P14: WT= 3.4 ± 1.1; Shank3^−/−^ = 2.7 ± 1.1; Welch’s t-test, p=0.1; adult: WT= 2.2 ± 0.8; Shank3^−/−^ = 2.1 ± 0.8; Mann Whitney test, p=0.7) but different decay time (in ms; mean ± sem; P14: WT= 16.8 ± 7.6; Shank3^−/−^ = 40.9 ± 19.9; Mann Whitney test, p=0.02; adult: WT= 22 ± 11.5, Mann Whitney test, p=0.03; Shank3^−/−^ = 30.2 ± 19, Mann Whitney test, p=0.03) (Figure 5B). Thus, mPFC efferents generate longer lasting synaptic currents in Shank3^−/−^ MD neurons resulting in an overall greater synaptic charge compared to littermate WT controls. The peak evoked synaptic current amplitudes did not differ between WT and Shank3^−/−^ mice (Figure S5B1-2), suggesting that the observed differences in EPSC decay rate were unlikely to be due to any differences in viral efficiency or spread from injection site.

**Figure 5.**
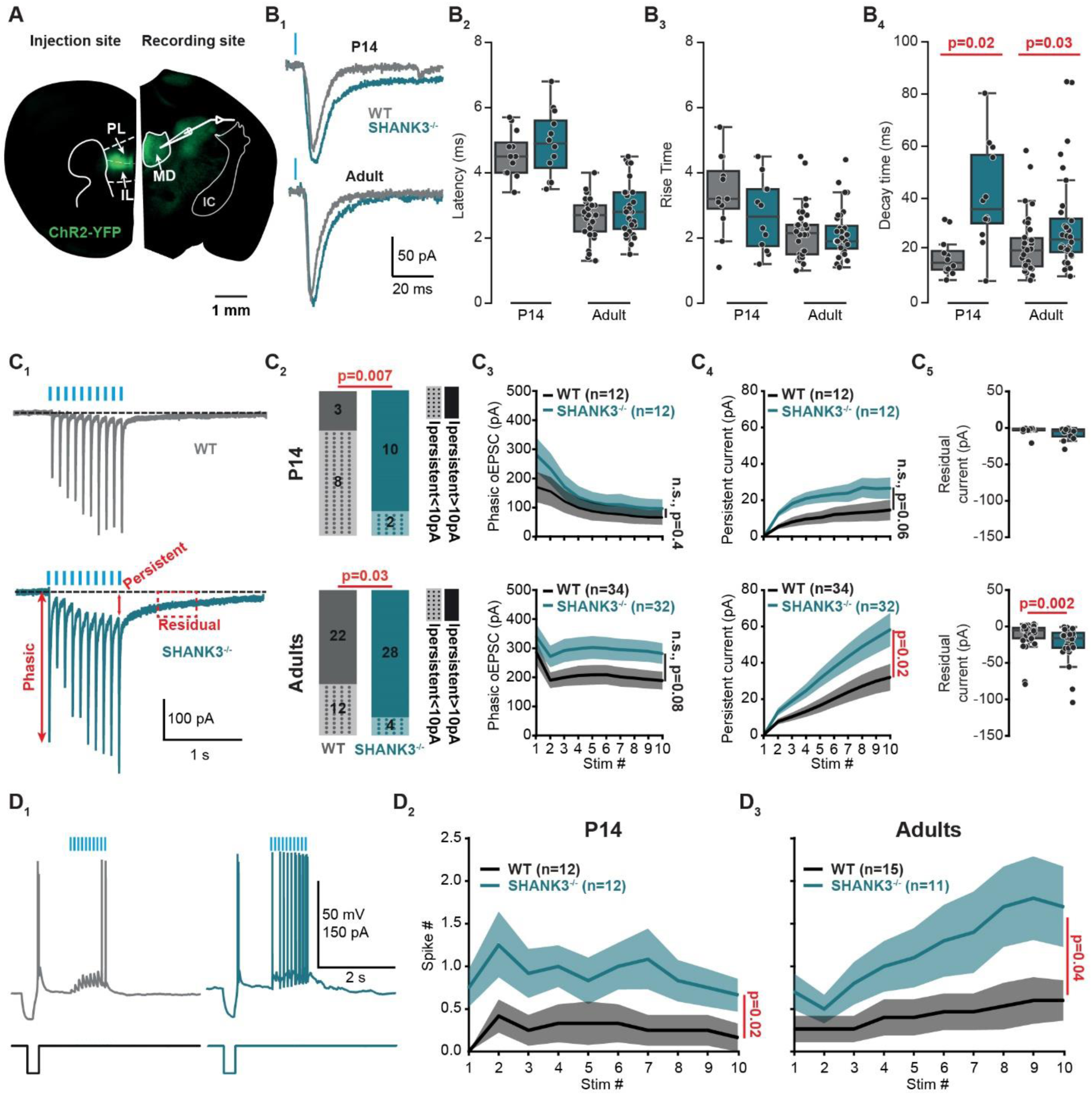
Enhanced mPFC-MD synaptic function in P14 and adult Shank3^−/−^ mice. (A) Epifluorescence image showing the injection site of AAV1_ChR2_YFP in mPFC (left) and labeled projections in MD thalamus (right). (B1) Representative optogenetically-evoked EPSC (oEPSC, phasic responses) from MD neurons of WT (grey) and Shank3^−/−^ (blue) mice. Data from P14 animals are shown on the top, while adult data are shown on the bottom. (B2-B4) Phasic oEPSC latency, rise time, and decay time for P14 (left) WT (grey, 2 mice, 12 cells) and Shank3^−/−^ (blue, 2 mice, 11 cells), and for adults (right) WT (grey, 16 mice, 34 cells) and Shank3^−/−^ (blue, 12 mice, 32 cells). (C1) Representative oEPSC trains elicited by 10 Hz stimulation. Phasic, persistent, and residual current measurements are indicated in red. (C2-5) P14 (top), Adult (bottom). (C2) Proportions of cells with detectable (>10 pA) or undetectable (<10 pA) persistent current (_Ipersistent_) in MD cells of WT (grey) and Shank3^−/−^ (blue) mice. (C3) Averaged phasic current amplitude with 10 Hz stimulation for WT (grey) and Shank3^−/−^ (blue) mice. (C4) Averaged persistent current with 10 Hz stimulation for WT (grey) and Shank3^−/−^ (blue) mice. (C5) Residual responses in WT (grey) and Shank3^−/−^ (blue) mice. (D1) Representative traces of current-clamp recordings from MD cells upon repeated activation of mPFC axons. (D2) Number of evoked action potentials during oEPSC trains from P14 WT (black) and Shank3^−/−^ (blue) mice. (D3) Evoked action potentials from adult WT (black) and Shank3^−/−^ (blue) mice. Traces represent mean ± SEM.

We noticed that during 10 Hz trains, the synaptic current did not fully decay to baseline between individual responses of the train (Figure 5C). We refer to this build-up as the persistent part of the synaptic response, which accumulates over the course of the oEPSC train. We classified MD neurons regarding whether they showed a persistent response based on the amplitude of the persistent current at the end of the oEPSC train (just before oEPSC #10, Figure 5C1) greater than 10 pA. In Shank3^−/−^ mice both the incidence and amplitude and of the persistent synaptic current showed a developmental progression. At P14 only 3/11 WT cells exhibited a persistent current, compared to 10/12 Shank3^−/−^ cells (Figure 5C2, top, chi-square test, p=0.007). In adult mice, persistent synaptic responses became more common even in WT mice, appearing in about 2/3 of MD cells (22/34 cells), whereas almost all Shank3^−/−^ MD cells exhibited a persistent current (28/32 cells, Figure 5C2, bottom, chi-square test, p=0.03). These results demonstrate a significant difference in the proportion of MD cells showing a late synaptic response between WT and Shank3^−/−^ mice at both P14 and adulthood. In adulthood, the persistent current build-up become greater in Shank3^−/−^ mice than in WT mice (Figure 5C4, bottom; WT vs. Shank3^−/−^, 2-way ANOVA with repeated measures: genotype p=0.02; stimulation p<0.0001; interaction p<0.0001).

We found that upon termination of oEPSC trains, MD neurons additionally displayed a slowly-decaying “residual” response lasting for several seconds, much longer than the decay time constant of individual oEPSCs reported above (<50 ms). We quantified the residual current as that measured 500 ms following the end of the stimulus train and found that it was greater in Shank3^−/−^ mice, but only in adults (Figure 5C5; mean ± sem; P14: WT=-3.5 ± 5.6; Shank3–/– = −7.9 ± 8.8; Mann-Whitney test, p=0.1; adult: WT = −12.2 ± 3.2; Shank3^−/−^ = −22.8 ± 4.1, Mann-Whitney test, p=0.002).

These findings indicate increases in connection strength between mPFC and MD neurons in Shank3^−/−^ mice. This synaptic strengthening becomes especially evident during repetitive activation with 10 Hz stimulus trains. This then triggers to greater recruitment of MD cells as measured by numbers of evoked action potentials (Figure 5D1-3, WT vs Shank3^−/−^, 2-way ANOVA with repeated measures, P14: genotype p=0.02; stimulation p=0.2; interaction p=0.9; adult: genotype p=0.049, stimulation p<0.0001, interaction p=0.0001).

### Synaptic receptors underlying the augmented function of the mPFC-MD synapse in Shank3^−/−^ mice

To investigate the nature of the persistent synaptic current in Shank3^−/−^ mice, we conducted a set of separate experiments using pharmacological compounds targeting two major receptor classes known to be affected by Shank3 mutations in other brain regions. Additionally, we chose to target receptors with a long duration of action, based on the previously observed increase in oEPSC decay time and the presence of late persistent current in Shank3^−/−^ mice (Figure 5). First, we used AP5, an antagonist of NMDA receptors. Second, we conducted an occlusion experiment using DHPG, an agonist of mGluR1 and mGluR5 receptors. These drugs were applied only if the recorded cell exhibited at least 10 pA of persistent current.

We first hypothesized that the increase in persistent current is mediated by NMDARs. Accordingly, bath application of AP5 should induce little to no reduction in WT mice, as oEPSCs recorded at −70 mV are predominantly mediated through AMPARs due to the magnesium block on NMDARs. Consistent with this, 0.1uM AP5 had no significant effect on phasic or persistent currents in WT mice. However, in Shank3^−/−^ mice, AP5 reduced both phasic (Figure 6 A, B; 2 way-repeated ANOVA, bath vs AP5, drug p=0.009, stimulation p=0.3, interaction p=0.9) and persistent current (Figure 6 A, C; 2 way-repeated ANOVA, bath vs AP5, drug p=0.0001, stimulation p<0.0001, interaction p<0.0001), indicating that the Shank3^−/−^ mutation results in abnormal functional activation of NMDARs.

**Figure 6.**
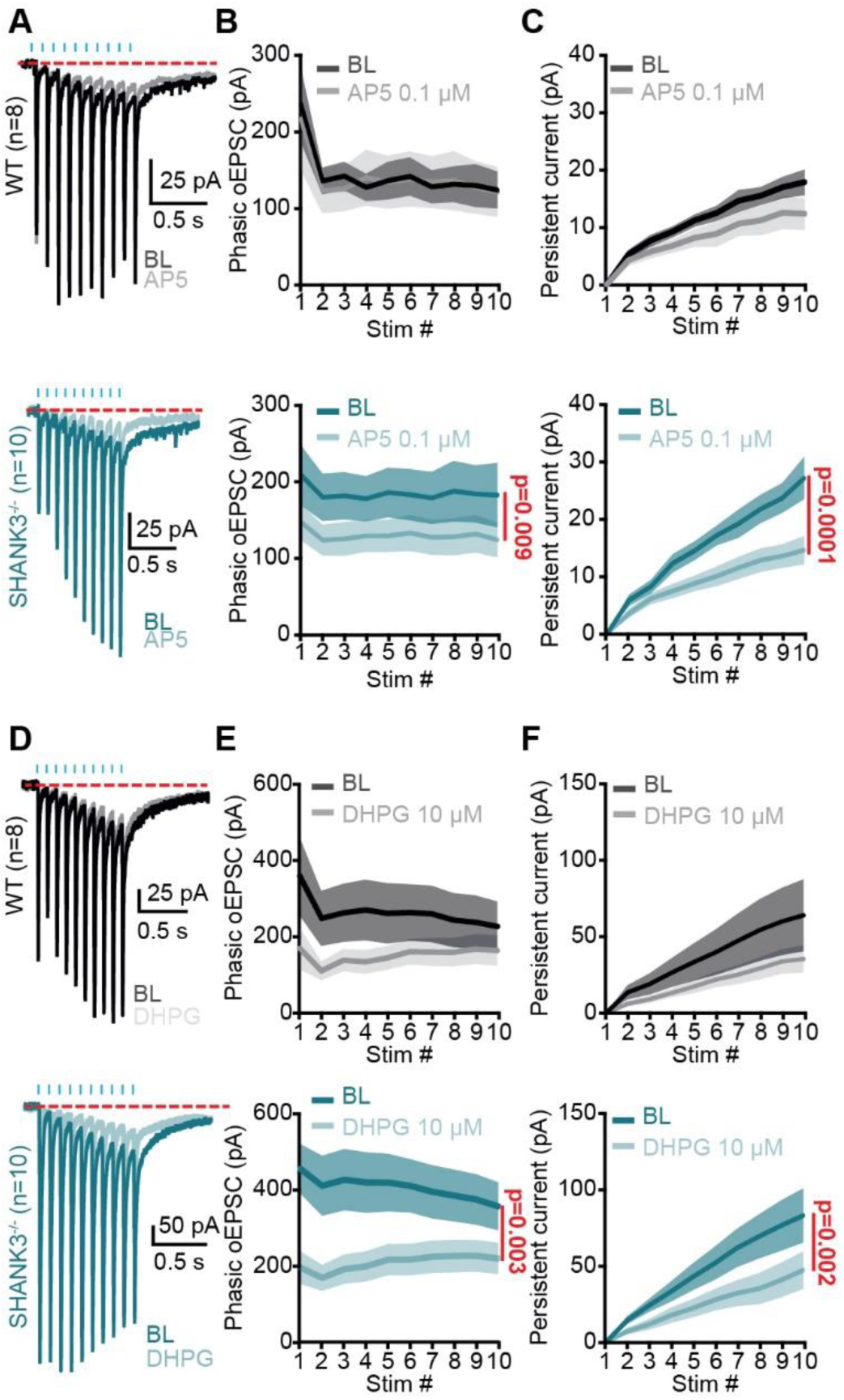
mGluR and NMDAR contribution to the phasic and persistent mPFC response to oEPSCs especially in Shank3^−/−^ mice. (A) Representative oEPSCs elicited with 10Hz stimulation in MD before (dark color) and after 0.1uM AP5 (light color) for WT (grey, 6 mice, 8 cells) and Shank3^−/−^ (blue, 3 mice, 10 cells) mice. (B) Peak amplitude of phasic currents in response to 10Hz stimulation in WT (top, grey) and Shank3^−/−^ (bottom, blue) mice before (BL, baseline, dark color) and during AP5 bath application (AP5 0.1uM, light color). (C) Same as (B) for persistent current. (D) Representative oEPSCs elicited with 10Hz stimulation in MD before (dark color) and after 10uM DHPG (light color) for MD recordings from WT (grey, 3 mice, 8 cells) and Shank3^−/−^ (blue, 4 mice, 12 cells) mice. (E) MD phasic current with 10Hz stimulation for WT (top, grey) and Shank3^−/−^ (bottom, blue) mice before (BL, baseline, dark color) and during AP5 bath application (AP5 0.1uM, light color). (F) Same as (E) for persistent current. Traces are represented are mean (solid line) ± sem (shaded line).

Our second hypothesis, based on the original Shank3^−/−^ complete knockout model(11), is that there is a dysregulation of postsynaptic mGluR5 signaling, which might be expected given that mGluRs show long-lasting responses(39). We initially tried to use MTEP, the antagonist of mGluR5, but the results were inconclusive. To further investigate, we performed an occlusion experiment using DHPG, a mGluR1 and mGluR5 agonist, reasoning that this would reduce the amplitude of the synaptic response, as a subset of post-synaptic mGluRs would already be activated by bound DHPG. We observed a decrease in both persistent (Figure 6D, F; 2-way repeated ANOVA, bath vs. DHPG, drug p=0.003, stimulation p=0.7, interaction p<0.0001) and phasic currents (Figure 6D, E; 2-way repeated ANOVA, bath vs. DHPG, drug p=0.003, stimulation p=0.7, interaction p<0.0001) in Shank3^−/−^ mice, but no change in WT mice. The lack of effect in WT mice suggests that mGluR1/5 signaling is tightly regulated and not recruited at either pre- or postsynaptic sites under normal conditions. In contrast, the reduction in phasic and persistent currents in Shank3^−/−^ mice upon DHPG application likely reflects a dysregulation of mGluR1/5 signaling.

## Discussion

In this study, we investigated the pathophysiological development of the medial prefrontal cortex (mPFC) and its output to the mediodorsal thalamus using the Shank3^−/−^ mouse model of autism spectrum disorder (ASD). After initially assessing function at P11, we mainly compared relevant developmental time points, specifically P14 (pre-weaning stage) and adulthood (>P55). Initially, we confirmed known ASD-related behavioral deficits, including impaired social behaviors in adult mice and altered ultrasonic vocalizations (USVs) in juvenile mice. We then employed our in vitro local field potential (LFP) recording design, which allowed us to identify a significant difference between the P14 and adult stages. At P14, we observed hypoexcitability, with a switch to mPFC circuit hyperfunction in adults. Furthermore, using whole-cell patch-clamp recordings of pyramidal cells, we uncovered distinct developmental patterns between cortical layers: at P14, layer 2/3 pyramidal cells exhibited hypoexcitability, while layer 5 cells showed signs of hyperexcitability. As the mice matured, the hyperexcitability in layer 5 cells was exacerbated, whereas layer 2/3 cells tended to normalize their excitability. Interestingly, we observed that the excitation/inhibition (E/I) ratio was altered specifically in layer 5 neurons at both developmental stages, while no significant changes were observed in layer 2/3 cells. These findings led us to further investigate the mPFC-mediodorsal (MD) thalamus pathway, a major connection dependent on layer 5 output and involved in social behavior. We observed significant alterations in this pathway, with changes in synaptic strength and post-synaptic receptor composition in Shank3^−/−^ mice, which were linked to disruptions in NMDA and mGluR group 1 receptor signaling.

### Longitudinal approaches in ASD research

While much ASD research focuses on mature animals, longitudinal studies are critical for understanding how genetic mutations influence neurodevelopment. Our study highlights the progression of deficits in Shank3^−/−^ mice, starting with early behavioral alterations at P11 and cellular hypoexcitability at P14, which then evolves into circuit hyperfunction in adulthood. The observed shift in mPFC network activity across development, from hypoexcitability to hyperfunction, is to our knowledge a completely novel finding. Observations of early cellular deficits have been reported in other studies using a different Shank3 mutant model, Shank3B^−/−^, in the corticostriatal circuit, where early abnormal hyperactivity of corticostriatal afferents was observed(40), however the long-term changes into adulthood were not reported. This group also confirmed early behavioral alterations in the Shank3B^−/−^ model(41). Altered USVs are also commonly observed in various genetic mouse models of ASD(10). These observations suggest that longitudinal approaches, such as our work, can provide critical insights into how gene mutations influence the progression of ASD over time and might help reveal potential targets and windows for therapeutic intervention.

### Layer-Specific mPFC Excitability in ASD

As mentioned above we found layer specific mPFC deficits that are different at P14 compared to the adults. The scientific literature regarding mPFC dysfunction has largely focused on mature animals(5,23,42,43). The hyperexcitability observed in mPFC layer 5 pyramidal cells is consistent with findings in other mouse model(5,35,44). However, other reports suggest that mPFC layer 2/3 cells can be altered in genetic mouse ASD models(30,43). Our intracellular recording data (Figure 3) revealed that while the intrinsic properties of layer 2/3 pyramidal cells are comparable between adult Shank3^−/−^ and WT mice, layer 5 pyramidal cells exhibit hyperexcitability in Shank3^−/−^ mice. A novel finding of our study is that these properties appear to be dynamic and development-dependent. Specifically, we detected hypoexcitability in layer 2/3 cells at P14, while layer 5 cells exhibited only a mild hyperexcitability—characterized by a depolarized resting membrane potential—at this stage. We speculate that compensatory mechanisms may normalize layer 2/3 cell excitability by adulthood, while at the same time this could, through enhanced output, lead to hyperexcitability in layer 5 cells in adult mice. Pyramidal cell morphology and intrinsic membrane properties develop in parallel in layers 2/3 and 5; however, the development of synaptic inputs differs between these layers(45) : during the first two postnatal weeks, layer 2/3 primarily receives excitatory inputs, while layer 5 does not(46). Shank3 is a scaffolding protein found in excitatory synapses(47,48). Its knockout may first affect early-developing synapses in layer 2/3 before impacting those in layer 5. This suggests that different layers of the mPFC may be affected differently to different ASD-related gene mutations.

Taking advantage of simultaneous recordings across layers with our in vitro multi-electrode array assay (Figure 2), we recorded LFP/CSD signals reflecting synaptic activity in mPFC neurons. Comparable to our patch-clamp data, we observed an interesting dynamic of signal excitability depending on age. At P14, we discovered that only the presynaptic signal (fiber volley) in layer 5 is hypoexcitable. This signal is likely dominated by thalamic inputs to the mPFC. In adulthood, we observed a global presynaptic hyperfunction of the mPFC across both layers, with layer 5 postsynaptic signals also hyperfunctional. Our results highlight the heterogeneity of ASD-related neurodevelopmental changes, where distinct mPFC layers and synaptic circuits exhibit alterations at different developmental stages. Consequently, these findings underscore the importance of considering both layer-specific and age-dependent parameters when studying the pathophysiological mechanisms linked to ASD.

### E/I ratio theory in ASD

Twenty years ago, Rubenstein and Merzenich hypothesized that ASD dysfunction may generally be related to increase in excitatory/ inhibitory ratio(38). This theory has been of interest of many(8,49) as it can relate to different ASD mouse models. Recently, Antoine et al. (50) has observed an elevated E-I ratio is a common circuit phenotype in four different genetic mouse models of ASD. They demonstrated that this elevation reflects stabilization of synaptic drive rather than driving network hyperexcitability, as was previously believed. In contrast, in our study (Figure 4), we observed a reduction in the E/I ratio in Shank3^−/−^ mPFC layer 5 cells at both P14 and in adults. Despite this, LFP recordings revealed global mPFC hyperfunction and layer 5 pyramidal neuron hyperexcitability in adults, though less so at P14. These results do not directly contradict the findings of Antoine et al. but suggest that the E/I ratio imbalance as observed at the single cell level may not necessarily be the cause of the observed network excitability changes. Instead, the reduced E/I ratio we observed may reflect an adaptive response to early cellular-level disruptions, possibly originating from developmental imbalances that influence network function later. Additionally, at this point we cannot exclude potential changes in interneuron excitability, as Shank3^−/−^ models have been shown to exhibit deficits in interneuron function, which could also contribute to the observed network alterations(51,52).

### mPFC-MD synaptic connectivity

Both P14 and adult Shank3^−/−^ mice exhibit altered mPFC-MD thalamus synaptic connectivity which results in a functional increase in mPFC-driven MD neuron spikes (Figure 5). Additionally, we observed a greater incidence of persistent and residual synaptic responses in MD neurons of adult Shank3^−/−^ mice which may contribute to behavioral dysfunction. In agreement with our observation of altered corticothalamic connectivity, it has been shown that sensory cortico-thalamic synapses are also altered in adults Shank3^−/−^ mice(53), also displaying a buildup of current under a 40Hz train stimulation.

Clues for a critical period of development for the mPFC-MD circuit have emerged. Indeed, anatomical lesions(54) or chemogenetic modulation to disturb the mPFC-MD circuit before adulthood results in physiological changes of mPFC activity(55,56) or social behavior(55). Similar modulation in adulthood failed to affect mPFC circuits or social behavioral changes. It is interesting to note that the density of connections between the mPFC and MD regions decreases at postnatal day 14 (P14)(12) and that a similar decrease in the expression of the SHANK3 transcript occurs at the same time point(18). Our findings emphasize significant alterations in mPFC-MD synaptic connectivity in Shank3^−/−^ mice, with functional changes evident across both juvenile and adult stages. The increased incidence of persistent synaptic responses in adult mice underscores the lasting impact of these early disruptions, highlighting the need to further understand how such changes contribute to behavioral dysfunction.

Importantly, the choice of NMDA and mGluR pharmacological manipulations was motivated by the longer decay times observed in MD cells of Shank3^−/−^ mice (Figure 6). This observation led to several mechanistic hypotheses, including receptor mislocalization(11), spillover effects(57), altered receptor composition(58,59), or an incomplete developmental switch(60). Interestingly, pharmacological modulation affected the persistent component in Shank3^−/−^ mice but had no detectable impact on WT, suggesting that NMDARs and mGluRs are indeed potential targets for this buildup of current in Shank3^−/−^ mice. In our study, we observed a novel population difference in the presence of persistent currents between Shank3^−/−^ and WT mice. Specifically, at both P14 and in adults, a majority of cells with from Shank3 mutants showed persistent currents, whereas WT mice displayed a higher proportion of cells with persistent currents at P14 (70%) and a reduced proportion in adults (35%) (Figure 5). This finding is consistent with the known alterations in NMDA receptor-mediated neurotransmission in other Shank3 loss of function models(55,61), where NMDA signaling is disrupted. Additionally, the occlusion experiment with DHPG provided further insight into the altered receptor signaling in Shank3^−/−^ mice. DHPG application led to also to a reduction in phasic current, suggesting that presynaptic alterations could contribute to the phenotype. Collectively, these findings reinforce the notion that Shank3 deficiency leads to disruptions in both NMDA and mGluR1/5 signaling pathways, ultimately affecting synaptic transmission.

### Caveats and limitations

Here we demonstrated novel findings of developmental ASD-related circuit dysfunction in a model of severe ASD, Shank3^−/−^ mice, at two important time points: P14 and adults (>P55). However, a potential limitation of our results is that they were mainly obtained in isolated brain slices. These have the advantage of being accessible for mechanistic and pharmacological characterization, using LFP or cellular physiology. However, the conditions *in vitro* can never completely match those in the intact brain, and any disease-related differences in extrinsic connectivity, neuromodulatory state, etc. will not be directly revealed by the in vitro approach. Yet the use of in vitro optogenetics and pharmacology has the advantage of leading to better understanding of cellular/molecular pathophysiological mechanisms directly related to genetic mutations. One additional potential drawback of LFP network analysis is that the mode of synaptic stimulation is through single extracellular stimulus pulse that evokes a simplified and highly synchronized input that may not model normal ongoing circuit activity in behaving animals. Yet being able to reliably and quantitatively determine input-output relationships provides an integrated and robust assay of circuit excitability and E/I ratio that together outweigh this limitation. Further, given that both WT and Shank3^−/−^ slices received comparable stimulation, yet displayed different responses, with hyperfunction of mPFC observed in adult Shank3^−/−^, it indicates that the circuit state in Shank3^−/−^ is more sensitive to synaptic input than in the WT. It would be interesting in the future to perform similar experiments *in vivo* by for example optogenetically stimulating the MD thalamus axon in mPFC while recording mPFC activity with linear silicon multielectrode probes. Another limitation of our study is that as of now, there is no direct evidence of a link between early circuit dysfunction and later behavioral effects. Future studies, using in vivo recording, like fiberphotometry(62) or miniscope(25) during the social interaction task will allow to relate the activity of mPFC (bulk fiberphotometry signal) or mPFC cells (miniscope) to the deficits. Finally, our study can be a starting point for establishing a causal link between early neurodevelopmental events and ASD-relevant neural circuit alterations. Acute mPFC circuit manipulation using optogenetics(62) or chemogenetics(55,63) in adults has been shown to be able to rescue some social behavior. It will be interesting to see if similar manipulations performed chronically during early life will have a significant impact on adult physiology and behavior. By identifying these critical periods of mPFC development, we might be able to perform early interventions that may prevent or mitigate ASD-related circuit dysfunction.

### Conclusions and future directions

This study uncovers a critical and never described shift in mPFC network activity in Shank3^−/−^ mice with distinct layer-specific alterations. Our findings highlight the progressive nature of ASD-related cortical dysfunction, emphasizing disruptions in the mPFC-mediodorsal thalamus pathway as a potential link to behavioral deficits. Crucially, this research underscores the importance of developmental timing in ASD, suggesting that early circuit dysfunction may set the stage for long-term impairments. By identifying these critical windows, our work lays the foundation for targeted early interventions, opening new possibilities for mitigating ASD-related neurodevelopmental disruptions.

## Materials and Methods

### Animals

Mice were bred and maintained according to Stanford University, School of Medicine Animal Research Requirements, and all procedures were approved by the Committees on Animal Research at Stanford University (protocol number: 12363). Mice carrying the *Shank3* complete knock out (Δe4–22) were a kind gift from Dr. Yong-Hui Jiang(11), Duke University. All comparisons were made using both genders. Mice used for experiments were separated in two distinct age group: P14 (+/− ½ day) and adults (2-4 months). The mice were maintained on a 12 hr light/dark cycle. Food and water were available ad libitum.

### Genotyping of transgenic animals

Shank3^−/−^ and Shank3 WT mice were genotyped by PCR using the following primers: wild-type forward GTGCCACGATCTTCCTCTAAAC, wild-type reverse, AGCTGGAGCGAGATAAGTATGC, mutant forward TTGCATCTGGGACCTACTCC and mutant reverse AAAGCACTGACTCCTCTCTTGG.

### Behavioral tests

#### Ultrasonic vocalization recordings

Ultrasonic vocalizations are typically emitted by pups from birth to the weaning period and are related to communication behavior. In this study, we elicited ultrasonic vocalizations (USVs) by isolating the pups briefly from their dams and littermates(64). Specifically, one hour before the recordings, on postnatal day 11 (P11) the breeders were removed from the home cage. The home cage with the remaining litter was then moved to the testing room and placed on a heating pad for one hour to maintain the body temperature. Pups were then individually isolated. Each tested pup was placed in a testing cage within a sound-isolated, darkened box, with an UltraSoundGate 116Unb recording system (Avisoft Bioacoustics) positioned 15 to 20 cm above. The USVs were recorded with a sampling rate of 384,000 Hz in a 16-bit format (recording range: 0-250 kHz) for 10 minutes using the Avisoft-RECORDER USG/USGH software. Finally, the pup was returned to its home cage with its littermates. Once the entire litter had been tested, the home cage was returned to its original room, and the breeders were placed back in the cage. The recordings were next analyzed using the Avisoft-SASLab Pro software (version 5.3.2-14; Avisoft Bioacoustics). In recordings of pup vocalizations over a 10-minute period, the first 2 minutes are excluded from analysis as the calls may result from handling-related stress. Fourier transformation (512pt FFT, Frame=100%, Hamming window with 75% of time overlap) was applied to obtain the spectrograms (500Hz: Frequency resolution; 0.5ms: Time resolution). USV detection was performed using automatic whistle tracking in SASLab. Monotonic calls with a minimal duration of 5 ms and slope of −15 to 6kHz/ms were counted. Number of calls was tested for Normality using a Shapiro-Wilk test. Between-group comparison was performed using a non-parametric Mann Whitney test. Data are reported as mean+/− sem.

#### Social preference test

To measure sociability, mice were tested in a social interaction task. A clean box was partitioned into three zones. A camera was mounted directly above the box to record videos that were analyzed offline. Tested mice were habituated to the empty box for ten minutes, the first day. The next day, a pre-test phase was performed using two empty cages, one at each end of the box. The tested mouse had 10 minutes to explore. The test phase is performed 10 minutes later. A novel object was placed within a perforated cage at one end of the box and a conspecific is placed in a cage at the opposite end. The placement of the novel object and never-met mouse was randomly assigned. Using EzTrack(65), the videos were scored to determine mouse position, including time spent in regions of interest (ROIs) drawn around the cage containing either the novel object or the conspecific. After checking for normality, the interaction time between an inanimate object or conspecific was tested using a Welch’s t-test.

### Extracellular multielectrode recordings

Extracellular recordings were performed on coronal neocortical 400 µm thick slices from P14 or adults (>P55) Shank3^−/−^or Shank3 wild type (WT) containing the medial prefrontal cortex (mPFC). Slices were recorded in a humidified oxygenated interface chamber at 34°C and perfused at a rate of 2 ml/min with oxygenated artificial cerebrospinal fluid (aCSF). The aCSF contained (in mM): 10 glucose, 26 NaHCO_3_, 2.5 KCl, 1.25 NaHPO_4_, 1 MgSO_4_, 2 CaCl_2_, and 126 NaCl (300 mOsm).

A linear silicon multichannel probe (16 channels, 100 µm inter-electrode spacing, NeuroNexus Technologies) was positioned normal to the medial surface and spanning across all layers of the mPFC to record evoked local field potentials (LFPs). The silicon probe was linked to a pre-amplifier (PZ5-32, Tucker-Davis Technologies (TDT)), and recordings were made using an RZ5D processor (TDT). Data acquisition was conducted using the Real-time Processor Visual Design Studio (RPvdsEx) tool and a custom Python code. LFP data were acquired at 25 kHz and filtered between 0.1 and 500 Hz. A bipolar tungsten microelectrode (each wire, 50-100 kΩ, FHC) positioned in upper layer 5 was used to deliver a bipolar electrical stimulation (0.1ms duration, intensity loops from 100 to 800 µA, interval of 15s). In order to obtain a more reliable index of the location, direction, and magnitude of currents underlying network activity, we derived the current-source density (CSD) from raw LFP signals(26). Assuming a uniform extracellular resistivity, the CSD can be estimated as the second spatial derivative of the LFP. In our experiment, CSD was computed in mPFC by algebraic differentiation of the LFP at adjacent electrodes in the linear probe spaced by 100µm and assumed a uniform extracellular resistivity of 0.3 S/m(66). In addition, the following drugs, DNQX (AMPAR antagonist; 25uM) and CPPene (SDZ EAA 494; NMDA competitive antagonist; 1µM) were applied in the bath to identify the pre and post synaptic component of our signals. Relevant components were next identified based on their distinct shape and their amplitude was measured for each slice. Between-group comparison was performed using a 2-way repeated measure ANOVA. Data are reported as mean+/− sem.

### Electrophysiological whole-cell patch-clamp recordings in mPFC slices

Mice were deeply anesthetized with isoflurane (for P14 mice) or pentobarbital sodium (i.p., 50mg/kg; for adult mice). P14 mice were directly decapitated. Adult mice were perfused transcardially with ice-cold (4°C), oxygenated (95% O_2_ /5% CO_2_) sucrose slicing solution containing (in mM): 234 sucrose, 11 glucose, 26 NaHCO_3_, 2.5 KCl, 1.25 NaH_2_PO_4_, 10 MgSO_4_, and 0.5 CaCl_2_ (310 mOsm). Brains were quickly removed and 250 µm-thick coronal slices of mPFC cortex were cut in sucrose ACSF using a vibratome (VT1200; Leica). Slices were transferred to a holding chamber filled with standard ACSF bubbled with 95% O_2_ and 5% CO_2_. The composition of aCSF was (in mM): 10 glucose, 26 NaHCO_3_, 2.5 KCl, 1.25 NaHPO_4_, 1 MgSO_4_, 2 CaCl_2_, and 126 NaCl (300 mOsm). Slices were allowed to recover 30 minutes at 37°C and were then incubated at room temperature (20–25°C) until recording.

Individual slices were transferred to a recording chamber and continuously perfused at 2ml/min with oxygenated aCSF at room temperature. Patch pipettes (3-5 MΩ) were pulled from borosilicate glass (Sutter Instrument, CA, USA) on a micropipette puller (Model P1000, Sutter Instrument, CA, USA) and filled with an intracellular solution containing (in mM): K-gluconate, 144; MgCl_2_, 3; HEPES, 10; EGTA, 0.5 and 2 mg/ml biocytin, pH= 7.2, 280mOsm. Whole-cell patch-clamp recordings were obtained from layer 2/3 or layer 5 pyramidal cells selected based on their morphology and electrophysiological profile. Recordings were performed at room temperature (20–25°C) using a patch-clamp amplifier (Multiclamp 700B, Molecular Devices) connected to a Digidata 1440A interface board (Molecular Devices). Signals were amplified and collected using pClamp 10.7. Cells with an a stable access resistance of <25MΩ were analyzed. Resting membrane potential was measured immediately upon obtaining the whole-cell configuration, and only cells with a resting membrane potential more hyperpolarized than −50 mV were selected. Resting membrane potential was computed as the average of the sweeps were no current was injected. Membrane potentials were not corrected for junction potential. Cells were held at their resting membrane potential and V/I curves were constructed from a series of current steps in 10pA and 20pA increments. Input resistance was determined from the slope of the V/I curve in the linear region within 5 mV of the resting membrane potential. Rheobase was calculated as the minimal depolarizing current step triggering an action potential. A tungsten bipolar stimulating electrode was placed in mPFC layer 2-3 to evoke synaptic responses. Cells were held at −70mV to record excitatory synaptic current and with the same stimulus intensity at +10 mV to record the inhibitory synaptic current. The peak amplitude of these excitatory and inhibitory evoked responses was used to calculate the excitatory-inhibitory ratio. Results are given as mean ± standard error of the mean (sem). Between-group comparisons were performed using parametric test. A p-value below 0.05 was considered statistically significant.

After recordings, slices were fixed overnight at 4°C in 4% paraformaldehyde/ PBS and then washed in PBS. Slices were processed using Alexafluor 488-conjugated streptavidin (1:500, Invitrogen) to reveal biocytin. After a series of wash in PBS, slices were mounted on gelatin-coated slides in Fluoromount-G for confocal microscopy.

### Morphological analysis

Zeiss LSM900 laser scanning confocal microscope was used to image the mPFC layer 5 biocytin labelled neurons with 63x objectives and a 488nm laser. Images were processed using ImageJ (National Institutes of Health; http://rsbweb.nih.gov/ij/). Neuronal tracing and analysis were performed using the SNT toolbox(67).

### Viral injections

For adult experiments, the procedure for viral injection has been described in detail(68). Briefly, young adult mice were anesthetized using isoflurane (3% induction, 1.5% maintenance) and received carprofen 5 mg/kg intra peritoneal injection. Mice received bilateral injection in mPFC (in mm from bregma, AP 1.8, ML +/−0.5, DV −2.6). A volume of 500nl of virus encoding the ChR2 (AAV1-CaMKIIa-ChR2(H134R)_eYFP-WPRE-HGH, 10^12^ GC, 100 nl/min, Penn Vector Core, Addgene 26969P) was delivered to the injection sites.

For P14 recordings, P0 mice were anesthetized on ice. Mice received a bilateral injection in the frontal area (in mm from lambda, AP: 3.5, ML:+/−0.1, DV:-1.5). A volume from 100 to 200nl of virus encoding the ChR2 (AAVDJ-CaMKIIa-ChR2(H134R)_eYFP, 10^12^ GC, 100 nl/min, Gene Vector and Virus Core at Stanford University, catalog #36) was delivered to the injection sites.

### Optogenetics experiments

ChR2-expressing mPFC axons to MD thalamus neurons were activated using blue LEDs stimulations (Thorlabs M450LP2, 450 nm, duration 1 ms, maximal light intensity 13 mW, 19 mW/cm2) or blue laser (Laserglow Technologies, 473 nm, duration 1 ms, maximal light intensity 8 mW, 10.8 mW/cm2). The mPFC-MD thalamus synaptic connection was evaluated using 10 light stimulations at both 1Hz and 10Hz, with one train administered every 20 seconds, while maintaining voltage clamp at −70mV. The resultant synaptic responses were evaluated in voltage clamp to measure synaptic current and in current clamp to quantify the number of action potentials. Synaptic properties parameters were measured using custom MATLAB code.

To investigate the receptor composition underlying the abnormal persistent current observed in adult Shank3^−/−^ mutant mice, either DL-2-Amino-5-phosphonovaleric acid (AP5, Sigma, A5282) at a concentration of 100 µM (broad-spectrum NMDA receptor blocker) or (RS)-3,5-Dihydroxyphenylglycine (DHPG, Tocris, #0342) at a concentration of 10 µM (selective group I mGluR agonist) were applied for 10 minutes in the bath after obtaining a stable baseline recording. Partial washout was obtained in some cases. The impact of AP5 and DHPG on the 10Hz voltage-clamp synaptic response was quantified and compared to the baseline recordings. Statistical analysis, 2-way ANOVA with repeated measure, was performed to evaluate the significance of the effects observed.

## Author Contributions

Gabrielle Devienne: Data collection, Conceptualization, Software, Formal analysis, Investigation, Writing - Original Draft, Visualization, Funding acquisition; Gil Vantomme: Data collection, Software, Validation, Formal analysis, Investigation; John R Huguenard: Conceptualization, Software, Writing - Original Draft, Funding acquisition.

## Acknowledgments

We thank all past and current lab members for their valuable feedback throughout this project. We are laboratory equipment, and supplies. We also thank Laura Denardo and Rui Peixoto and their team for their invaluable advice on P0 virus injections.

