## SUPPLEMENTARY for "Disrupted Development of the mPFC-Thalamic Circuit in the Mouse Shank3^−/−^ model of Autism"

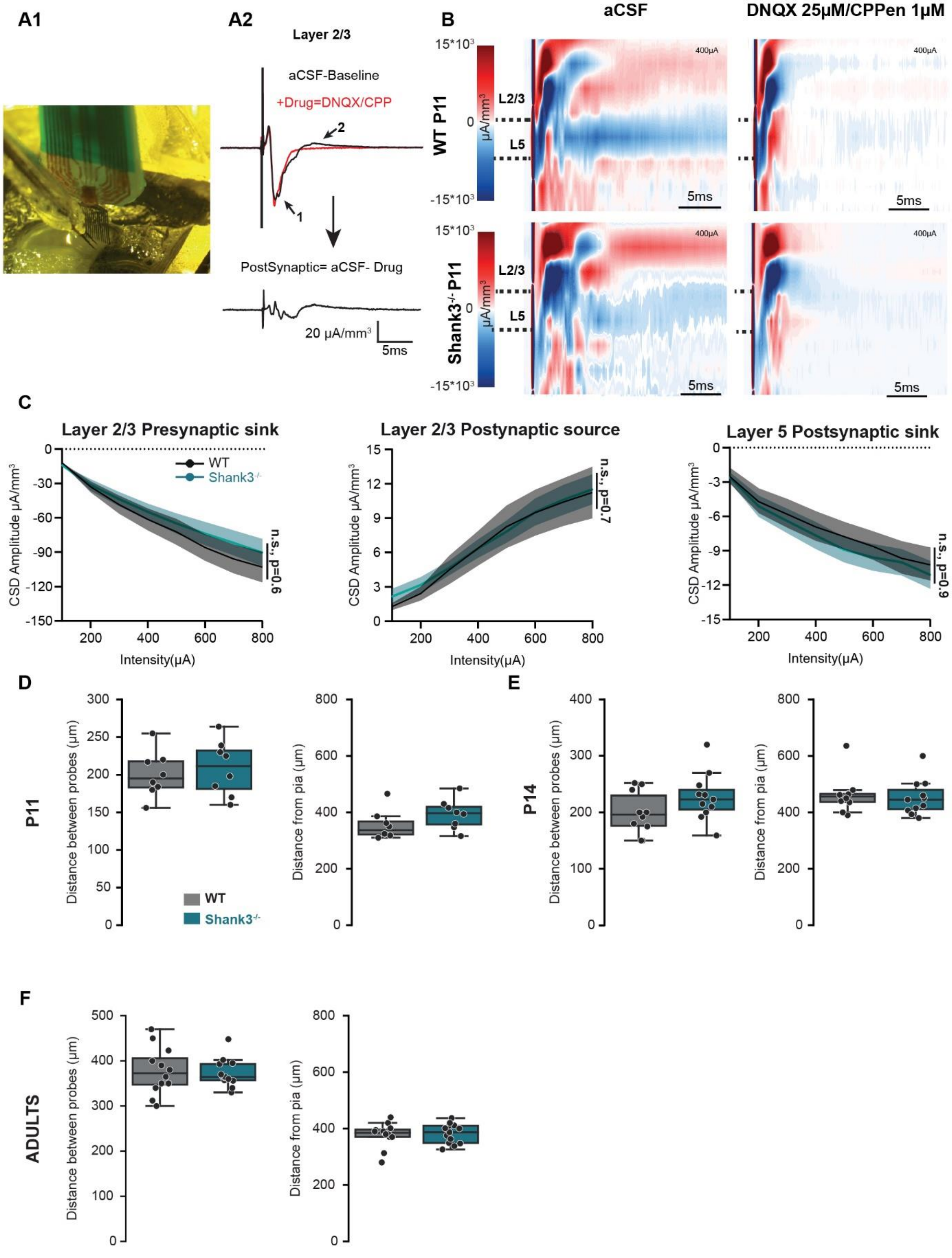


**Supplementary Figure 1. mPFC slices from Shank3$^{-/-}$ mice display normal excitability at P11.** (A1) Electrode placement for *in vitro* mPFC LFP recordings. (A2) Top panel: Representative CSD response recorded from the channel located in layer 2/3 under baseline conditions in ACSF (black). The trace exhibits two distinct components: a presynaptic component (indicated by arrow #1) and a postsynaptic component (indicated by arrow #2). Red trace: Following the application of DNQX (25 μM) and CPPEN (1 μM), only the presynaptic component (arrow #1) remains, as the postsynaptic component is blocked. Bottom panel: The subtraction of the drug-treated trace from the ACSF trace reveals the relatively smaller, isolated postsynaptic component. (B) Averaged CSD heatmap of mPFC network during 400 µA stimulation in P11 in WT (top) and Shank3$^{-/-}$ (bottom) littermates, before and after DNQX25uM +CPPene 1uM bath application (8 slices in each case, from 2 animals). Postsynaptic responses are blocked by DNQX/CPPene, revealing the short-duration pre-synaptic component that is essentially complete within 5ms of the stimulus. This presumably reflects action potential firing in presynaptic axons, as it was completely blocked by the sodium channel antagonist tetrodotoxin (not shown, n=3). Accordingly, responses blocked by DNQX/CPPene are postsynaptic, and mainly occur later than 5ms from stimulus onset. (C) Input/Output relationships for CSD peak amplitude across stimulus intensities ranging between 100 µA and 800 µA. Traces represent the mean (solid line) ± sem (shaded line) across multiple slices (n=8 slices, from 2 mice), and components 1, 2 and 3 are defined as in Figure 2. (D, E, F). Stimulation and recording electrode positions were equivalent for each genotype and at each developmental state: P11, P14 and adult, respectively. The distance between probes, i.e. the lateral distance between the stimulating electrode and the Neuronexus probe is shown on the left and the axial distance between the stimulating electrode and the pia is shown on the right. WT are in grey, Shank3$^{-/-}$ in blue.

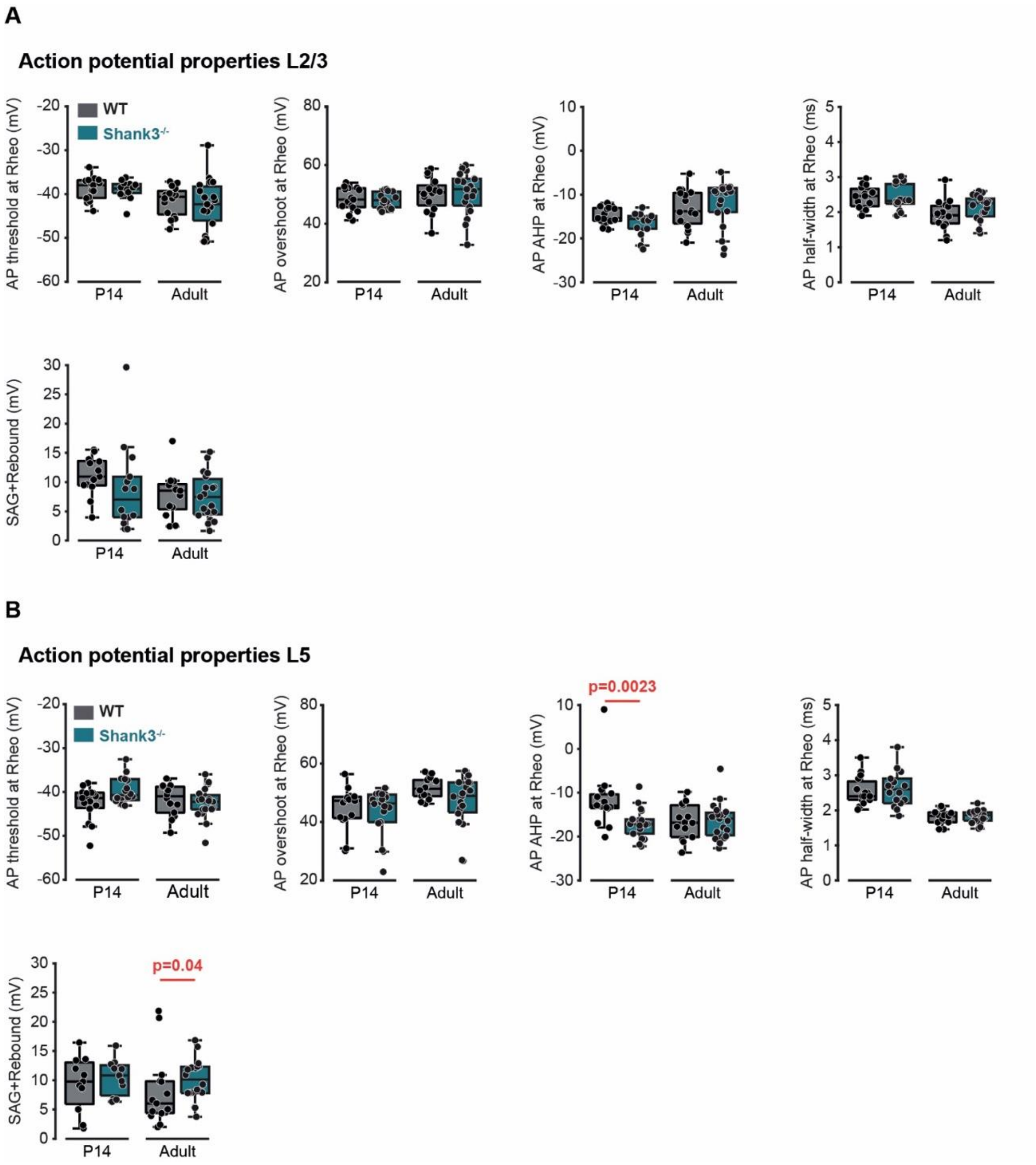


**Supplementary Figure 2. Layer 2/3 and layer 5 mPFC pyramidal cell action potential properties are largely unaffected in Shank3$^{-/-}$ mice, both at P14 and in adulthood.** (A) Quantification of mPFC layer 5 pyramidal cells rheobase (rheo) action potential properties (threshold, overshoot, afterhyperpolarization (AHP), half-width) and sag+rebound, at P14 and adults for WT (grey) and Shank3$^{-/-}$ (blue). Sag+rebound is a composite measure of hyperpolarization-induced membrane potential sag and overshoot (see methods). (B) Quantification of mPFC layer 5 pyramidal cells rheobase action potential properties at P14 and adults for WT (grey) and Shank3$^{-/-}$ (blue). L5 cells show differences in only two properties, action potential afterhyperpolarization, and sag+rebound.

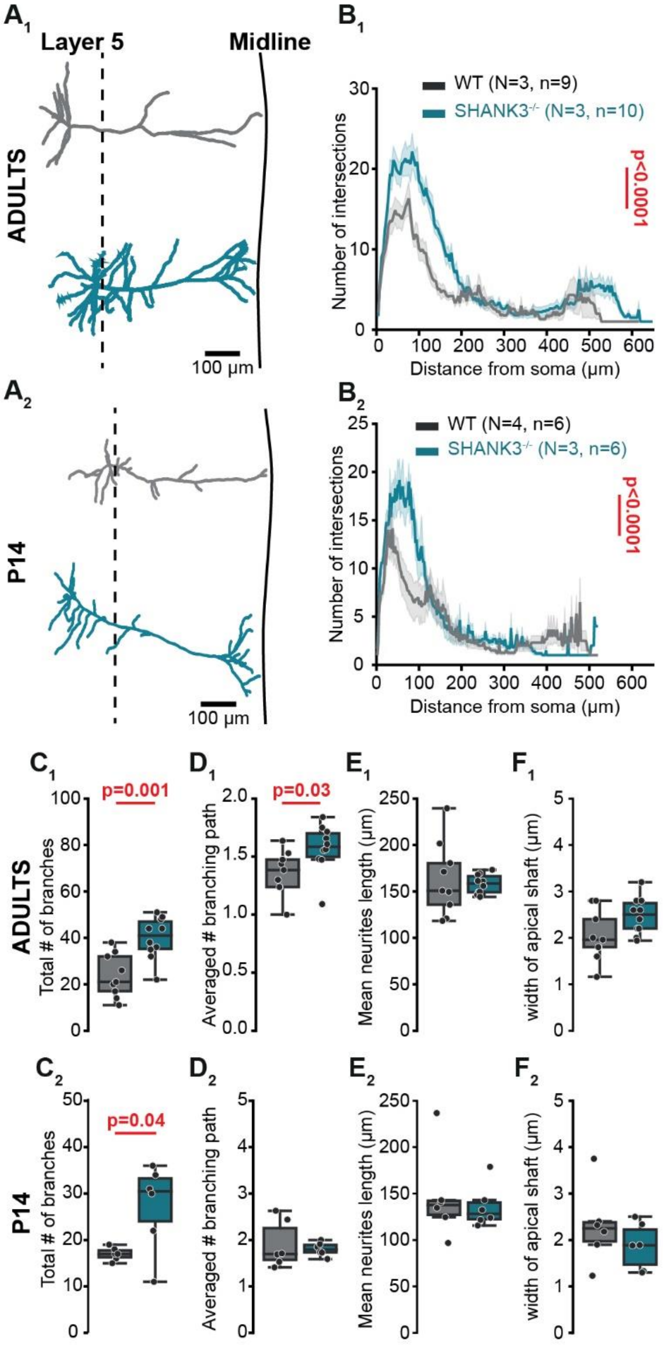


**Supplementary Figure 3. mPFC layer 5 pyramidal neurons display increased dendritic complexity in Shank3$^{-/-}$ mice both in early life and in adulthood.** (A) Representative examples of mPFC layer 5 pyramidal cells of adults (A1) and P14 (A2) WT (grey) and Shank3$^{-/-}$ (blue) mice reconstructed using biocytin filling after patch-clamp. (B) Average Scholl analysis of mPFC layer 5 pyramidal cells of adults (B1) and P14 (B2) WT (grey) and Shank3$^{-/-}$ (blue) adult mice. Traces are represented as mean ± sem. (C-F) Quantification of pyramidal cells morphological parameters (total number of branches, averaged number of branches, mean neurites length and width of the apical dendritic shaft) in adults (1) and P14 (2). WT are in grey, Shank3$^{-/-}$ in blue. (N is number of mice, n is the number of cells).

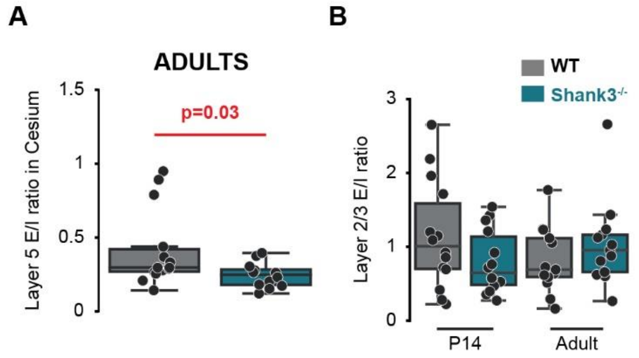


**Supplementary Figure 4. Alterations in excitatory-inhibitory ratio in mPFC layer 5 pyramidal cells but not in layer 2/3 pyramidal cells of Shank3$^{-/-}$ mice.** (A) E/I ratio of adults mPFC layer 5 pyramidal cells, as determined by the ratio of electrically evoked synaptic currents measured at -70 mV and +10 mV, with a cesium based intracellular solution. (B) E/I ratio of mPFC layer 2/3 pyramidal cells at P14 (left) and adults (right). WT are in grey, Shank3$^{-/-}$ in blue.

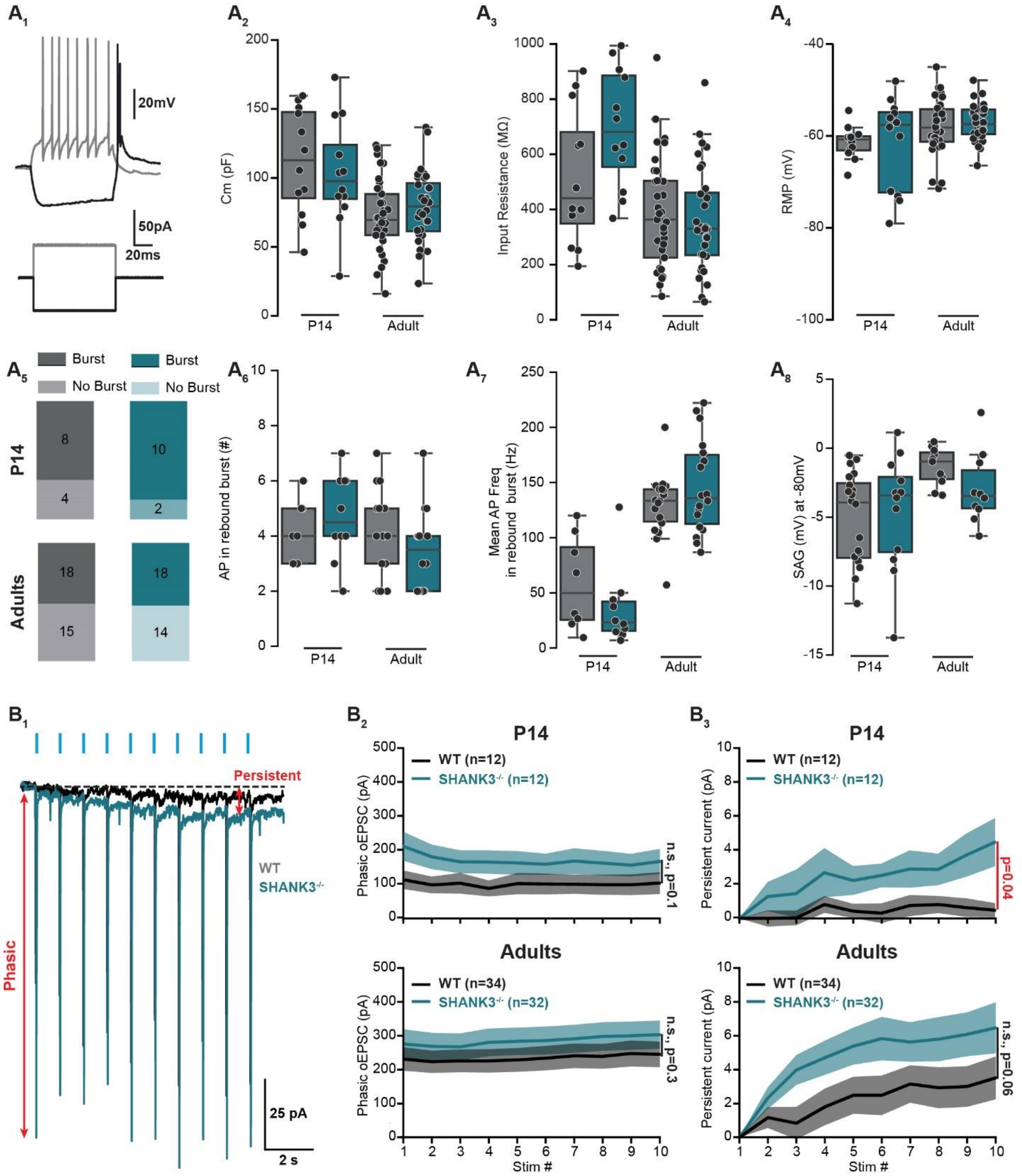


**Supplementary Figure 5. Intrinsic electrical properties of MD neurons are unaffected by Shank3 knockout. mPFC-dependent 1 Hz excitatory synaptic responses show phasic and persistent components.** (A) Example electrophysiological trace of an MD cell firing a rebound burst after a 50pA hyperpolarizing step (black) or tonic firing during a 50pA depolarizing step (grey) (1). Quantification of capacitance (Cm, 2), input resistance (3) and resting membrane potential (4) of adult MD cells of WT (in grey) and Shank3$^{-/-}$ (in blue). Characterization of the number of MD cells displaying a rebound burst (5), the number of action potentials within the burst (6), the mean action potential frequency within the burst (7), and the SAG measure when hyperpolarization reaches -80 mV in a subset of MD cells (8). (B) 1Hz optically evoked

EPSCs (oEPCSs) show phasic and persistent responses, evident during stimulus trains. (B1) Representative oEPSC elicited with 1Hz stimulation in MD. Phasic and persistent measurement are indicated in red. (B2) Averaged phasic current quantification at 1Hz stimulation across 10 stimulations for adult WT (grey) and $Shank3^{-/-}$ (blue) mice. (B3) Averaged persistent current quantification at 1Hz stimulation across 10 stimulations for adult WT (grey) and $Shank3^{-/-}$ (blue) mice.

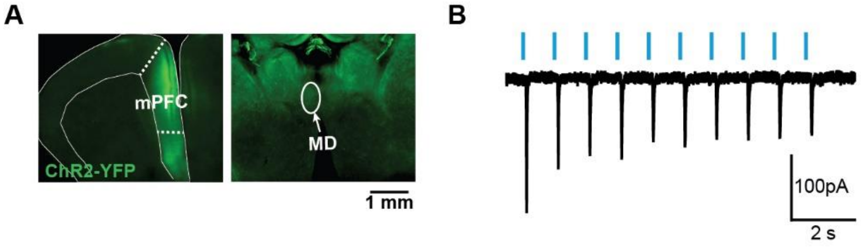


**Supplementary Figure 6. Early postnatal mPFC injections result in MD axonal labeling and functional synaptic responses.** (A) A labeled image showing the injection site in the mPFC at P14. This injection was performed at P0, with axonal labeling visible in the mediodorsal thalamus (MD). (B) An optogenetically evoked response recorded via patch clamp in a cell within the mPFC, indicating direct synaptic activation.
